# Literature-Based Discovery beyond the ABC paradigm: a contrastive approach

**DOI:** 10.1101/2021.09.22.461375

**Authors:** Erwan Moreau, Orla Hardiman, Mark Heverin, Declan O’Sullivan

## Abstract

Literature-Based Discovery (LBD) aims to help researchers to identify relations between concepts which are worthy of further investigation by text-mining the biomedical literature. The vast majority of the LBD research follows the ABC model: a relation (A,C) is a candidate for discovery if there is some intermediate concept B which is related to both A and C. The ABC model has been successful in applications where the search space is strongly constrained, but there is limited evidence about its usefulness when applied in a broader context.

Through a case study of 8 recent discoveries related to neurodegenerative diseases (NDs), we show the limitations of the ABC model in an open-ended context. The study emphasizes the impact of the choice of source data and extraction method on the resulting knowledge base: different “views” of the biomedical literature offer different levels of accuracy and coverage. We propose a novel contrastive approach which leverages these differences between “views” in order to target relations between concepts of interest. We explore various parameters and demonstrate the relevance of our approach through quantitative evaluation on the 8 target discoveries.

The source data used in this article are publicly available. The different parts of the software used to process the data are published under open-source license and provided with detailed instructions. The main code for this paper is available at https://github.com/erwanm/lbd-contrast (required dependencies are detailed in the documentation). A prototype of the system is also provided as an online exploration tool at brainmend.adaptcentre.ie.

## 1. Introduction

Nowadays virtually any biomedical research work is available almost instantly in digital form, but exploring the literature is made challenging by the ever-increasing amount of publications. Literature-Based Discovery (LBD) is an application of text mining which aims to automatically extract new insights from the scientific literature (Henry & McInnes, 2017). LBD aims to assist researchers in identifying potentially interesting relations between concepts, ideally contributing to faster and broader scientific progress.

In his seminal work Swanson (1986a) demonstrates that drawing links between two unconnected parts of the literature can lead to fruitful discoveries. The principle can be summarized as follows: given two subsets of literature respectively related to concepts A and C, retrieve the most important concepts associated with A and those associated with C. If the overlap B of these two sets is not empty, the B concepts are prime candidates for establishing a formal link between concepts A and C. The *closed discovery* variant of this approach assumes that both target concepts A and C are known from the outset, while in the *open discovery* variant one starts from a single target concept A.

This idea gave birth to the field of LBD, and the ABC model became the standard and virtually unique approach in the field (Henry & McInnes, 2017; Thilakaratne et al., 2019). The ABC model is both intuitively meaningful and technically convenient, as it is well suited for data representations such as knowledge graphs or vector-space models. Virtually every recent LBD work relies on the ABC model, for example (Lever et al., 2017; Pyysalo et al., 2019; Crichton et al., 2020). However, despite its advantages, the ABC model has some signficant limitations: first it is only intended to find a subset of potential “discoveries”, those which can be represented as two concepts which have a strong link through a third concept; it also tends to generate a quickly overwhelming number of relations. Arguably, this could explain why the ABC model led to to successful discoveries mostly in quite specific applications such as drug repurposing and pharmacovigilance Henry & McInnes (2017), which target very specific types of relations and concepts.

Smalheiser (2012) provides an insightful analysis of the subtle biases of the ABC model, and advocates for broadening the spectrum of methods used in LBD. Also in (Smalheiser, 2017) the same author emphasizes the original concept of “Undiscovered Public Knowledge” (UPK) (Swanson, 1986b), which forms the epistemological basis for the field. From this point of view, the ABC model can be seen as an implementation of UPK which has the merit of formalizing it in such a way that it can be exploited by automatic methods. Nevertheless it is “only one of several different types of models that can contribute to the development of the next generation of LBD tools” (Smalheiser, 2012). In the present work we propose such an alternative approach.

The studied literature can be represented in many ways depending on the options selected for extracting concepts from the documents: which initial set of documents is considered, which concepts are extracted, how concepts are filtered and counted, etc. These choices of representation are usually considered as fairly minor pre-processing options, even though they can have a significant impact on the resulting representation of the literature. In particular these choices can lead to different accuracy/coverage ratios, in turn causing different outcomes in the relations retrieved by the LBD system. We hypothesise that these different *views* of the literature (i.e. the representations obtained by different combinations of options) can be fruitfully combined for generating relevant candidate relations. In particular we explore a simple method where a *reference view* of the literature is contrasted with a *mask view* of the same literature in order to reveal relations which do not otherwise appear at the top of the ranking.

The contrastive method proposed in this paper is a simple parametric heuristic which does not rely on Machine Learning techniques. We purposefully adopt an unsophisticated approach for two reasons: as an exploratory attempt to reframe the problem of LBD, it makes sense to investigate simple methods first. This is also intended to preserve interpretability and transparency, in order to minimize the technological obstacles to adoption by biomedical experts.

The paper is structured as follows: in §2 we explain why state of the art LBD approaches are not suitable to retrieve certain types of relations, then we present some observations which form the basis of our approach. In §3 we detail our experimental methodology and introduce the LBD contrast method. The results of the validation experiments are discussed in §4, as well as a brief introduction to the online prototype.

## 2. Approach

### 2.1. Motivation

This work is motivated by the observation that a particular relation between two concepts is sometimes *known* in the sense that the relation is mentioned in the literature, yet it is *overlooked* in the sense that its significance has not been fully realized or investigated by the scientific community. This informal observation is supported by several cases in the domain of Neurodegenerative Diseases (NDs) which form the basis of this study. These eight relations are presented in Table 1 and share the following characteristics:

- The first co-occurrence is found before 1980 in 6 out of 8 cases, with two cases even found in the 50s. This is often followed by a few sporadic co-occurences across the years, without any visible surge in frequency during a long perid of time.
- Most cases show a strong revival at some point in the last two decades, marked by a significant surge in the number of co-occurrences.

**Table 1:** ND-related relations used as “overlooked discoveries” cases. ALS stands for Amyotrophic Lateral Sclerosis. The first co-occurrence year is taken from whichever data source contains the earliest co-occurrence.

| Concept 1 | Concept 2 | First Year |
| --- | --- | --- |
| ALS | Autism | 1990 |
| ALS | Extrapyramidal Disorders | 1967 |
| ALS | Spinocerebellar Ataxia | 1965 |
| Ataxia | ALS | 1979 |
| Frontotemporal Dementia | Bipolar Disorder | 2000 |
| Myopathy | ALS | 1954 |
| Parkinson Disease | ALS | 1951 |
| Psychotic Disorders | ALS | 1960 |

The recent surge in most of the cases is due to the fact that the body of knowledge about NDs has significantly improved recently thanks to various discoveries, in particular related to common genetic factors between diseases. For example, in 2011 the identification of C9orf72 repeat expansions as the most common genetic variant associated with both amyotrophic lateral sclerosis (ALS) and frontotemporal dementia (FTD) confirmed the previously recognized important pathobiologic association between these two conditions. In this case and in many others, the partial similarities between diseases were fairly well known but were not sufficiently clear to be considered as a significant discovery. Thus these cases are good examples of undiscovered public knowledge (Swanson, 1986b) worthy of being retrieved by LBD. However their properties make them unsuitable for discovery through the ABC model. The first problem is whether the target relation A-C exists or not, given that a few co-occurences are found before the time of the true discovery. The traditional ABC model assumes that either a relation exists or it does not, and its goal is to find relations which do not already exist. The second problem is the lack of a candidate B concept as link between A and C: in all these cases the only pre-existing indicators of a relation are some poorly characterized clinical symptoms in common. These form a much less clear-cut picture than the ideal scenario of the ABC model, which is meant to match fine-grained specific concepts with few connections rather than generic concepts related to many other concepts.

In fact, the prevalence of the ABC model may have introduced some unconscious biases towards evaluation methods and benchmarks discoveries which are compatible with it:

- On the one hand, *qualitative LBD* research, e.g. (Pyysalo et al., 2019), relies on a very small set of gold-standard discoveries, among which the initial LBD discoveries made by Swanson (1986a) are still the most widely used benchmark. While these relations represent real high quality discoveries they also tend to relate to very specific concepts, and this probably explains their rarity: in general discoveries often involve at least one quite generic concept, a case that the ABC model cannot handle very well.
- On the other hand, *quantitative LBD* research, e.g. (Lever et al., 2017), makes the simplifying assumption that any co-occurrence appearing at some point in time in the literature is a discovery. This assumption makes the use of advanced ML methods possible and evaluation more reliable from a statistical perspective. However this results in considering as discoveries an extremely wide range of relations, most of which an expert would discard as irrelevant. Thus while the quantitative approach is a reasonable simplification from a technical point of view for evaluation purposes, its usability in real applications is questionable.

The present work is an approach based on a small set of relations considered as relevant by ND experts. The ambition is to retrieve discoveries that researchers find useful, whether this is convenient or not from a technical point of view. Since state of the art ABC-based methods cannot handle these cases, we simply turn to an alternative approach. We do not propose a semantic model of discovery like the ABC model, instead we frame the problem as an agnostic data-driven task:^1^ given a set of validated discoveries, can a LBD system learn how to retrieve similar discoveries? This can be interpreted as a middle-ground between the qualitative and quantitavive LBD approaches: it does not impose restrictions on the gold-standard discoveries, nor does it indiscriminately assume that any co-occurrence is a valid discovery.

### 2.2. Data

Throughout this paper we use two distinct methods to extract the concepts and their frequency by document.^2^ This is meant to study the extent of the discrepancies between source datasets and to measure the impact of this choice on the results obtained by the LBD method proposed in this work. The raw source data is made of all the titles and abstracts extracted from Medline^3^ as well as the full-text articles from PubMed Central Open-Access (PMC-OA)^4^. The two datasets, denoted KD and PTC, are obtained as follows:

- **KD:** We use a modified version^5^ of the “Knowledge Discovery” code by Lever et al. (2017) to extract the concepts from the raw data sentence by sentence. The concepts are identified by string matching using the UMLS Metathesaurus^6^ as a list of target terms, filtering only the following UMLS semantic groups: Anatomy, Chemicals & Drugs, Disorders, Genes & Molecular Sequences and Physiology. An additional step of disambiguation is performed using an ad-hoc disambiguation system.^7^ The extracted concepts are represented as UMLS Concept Uniques Identifiers (CUIs).
- **PTC:** PubTatorCentral^8^ offers Medline and PMC contents conveniently annotated with concepts from several state-of-the-art text mining systems (Wei et al., 2019). The extracted concepts are represented as MeSH descriptors^9^

In this work we hypothesize that exploiting different representations of the literature can contribute to LBD. In this perspective we explore the difference between abstracts and full articles (as studied in (Westergaard et al., 2018)), this is why the data is decomposed into three different “views” for each data source:

- The *abstracts* view contains all the abstracts from Medline. Abstracts are expected to provide a narrow and focused picture of the literature, possibly missing some relations but faithfully reflecting the most important ones.
- The *articles* view contains all the full papers from PMC. Articles are expected to provide a large coverage and potentially more noisy representation of the literature.
- The *abstracts+articles* view contains the union of Medline abstracts and PMC full articles. The abstract of a paper which appears in PMC is discarded from Medline in order to avoid duplicates. This view is supposed to provide the most complete picture of the literature.

It is worth noting that the proportion of papers published as full article has drastically increased across the years, potentially causing a bias in the exploitation of the different views. Finally how concepts co-occurrences are counted is also a major parameter for the data representation: we use two variants where concepts are considered as a co-occurrence when they appear (1) in the same sentence or (2) in the same document. The former is assumed to provide fewer but more reliable matches since it is unlikely for two concepts to appear so closely by chance. With the latter one expects a larger coverage of the relations including more noise.

### 2.3. Preliminary observations

Fig. 1 shows the evolution of the frequency of co-occurrences across time for 7 of the relations in Table 1.^10^ While the two data sources tend to correlate across cases, important differences in frequency can be observed. In particular PTC shows no co-occurrence at all in the case of Bipolar-FTD. In fact PTC fails to identify the term “frontotemporal dementia” as the concept FTD, instead it identifies only the more general concept of “dementia”. On the contrary PTC appears to identify more co-occurrences in the case of ALS-Extrapyramidal Disorder. The frequency of co-occurrences is generally higher by document than by sentence since more pairs of concepts are matched in the former variant. However the latter can occcasionally be higher because several co-occurrences by sentence are counted in the same document. These differences illustrate the high variations due to the parameters guiding how the literature is represented.

**Figure 1:**
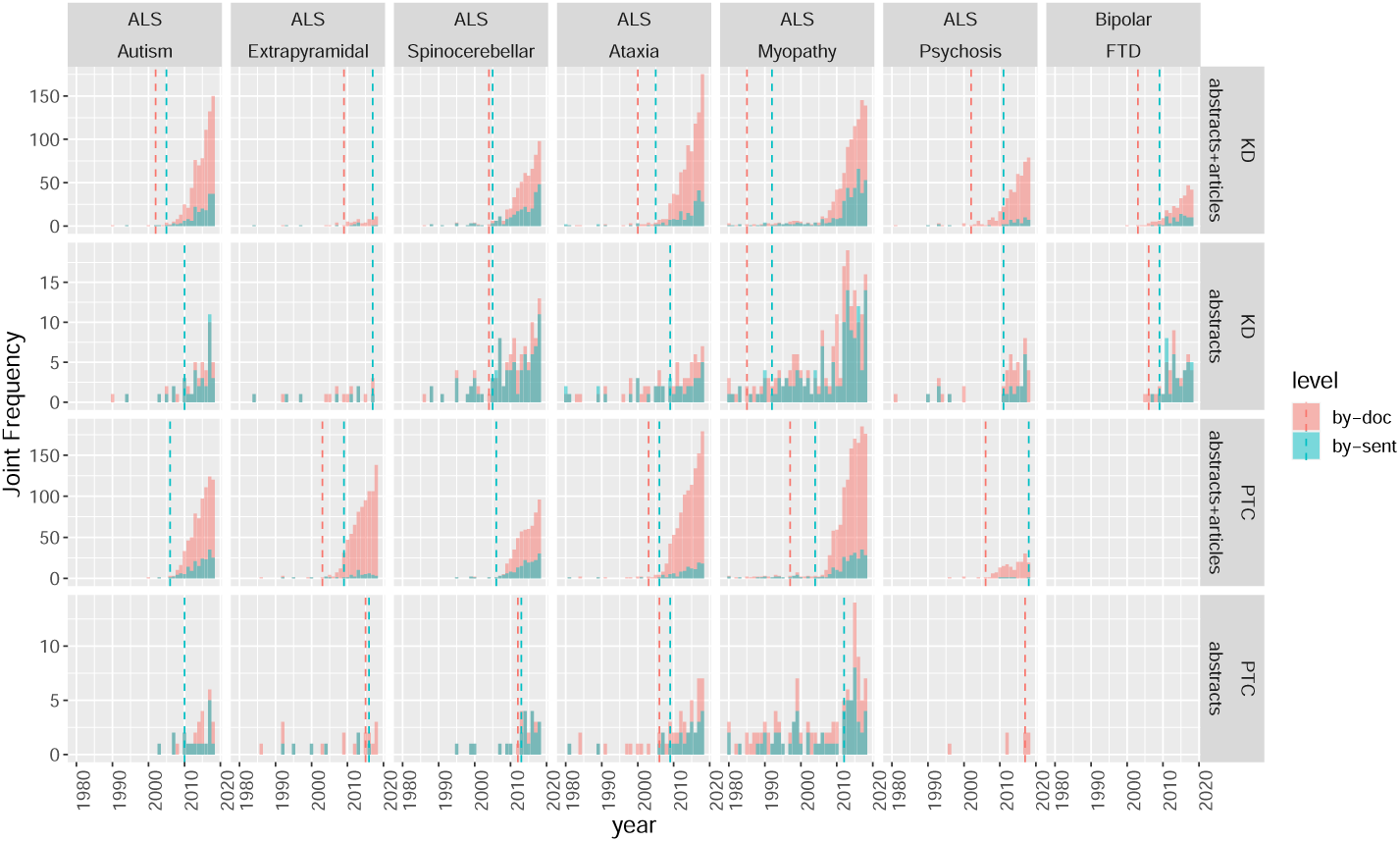
Co-occurrence frequency from 1980 to 2018 for 7 of the target relations, depending on data source and extraction method. The rows represent different data sources, while the colors represent the level at which co-occurrences are counted (by document or by sentence; see details in §2.2). The dashed vertical lines represent indicate the start of a consecutive non-zero count sequence of years.

As indicated in §2.1, the cases exhibit a similar pattern, albeit with some variations: a few sporadic co-occurrences in the 1980s and 1990s followed by a drastic surge in the 2000s. It is worth noting that the surge cannot be explained by the normal increase in number of publications across time, which is fairly regular around 4% by year for documents and around 6% for sentences. Naturally this surge pattern is due to researchers starting to study the relation more intensively due to its renewed interest/significance for the field. In this work we propose to define this surge as the marker of a discovery, and consequently the goal of a LBD system would be to retrieve relations susceptible of exhibiting such a surge in the future.

## 3. Methodology

### 3.1. Evaluation design

Our evaluation follows the methodology used in various other LBD works e.g. (Lever et al., 2017; Pyysalo et al., 2019). First we assume a target concept of interest; in our sample of ND-related discoveries (see Table 1), the target concept is the main disease, namely ALS (7 relations) or FTD (1 relation). The LBD system is provided with the target concept (e.g. ALS) and only the data from the years preceding the discovery year. It is expected to return a ranking of the concepts potentially related to the target, in which ideally the second concept of the relation (e.g. Autism) is ranked close to the top.

The cut-off year is meant to represent the last point in time before the discovery happens, as represented by the surge visible in Fig. 1. The choice of a cut-off year is made difficult by the various shapes of the frequency graphs, which differ depending on the relation, the data source and the level considered. Interestingly, this difficulty illustrates that the notion of discovery is often blurry, even though the traditional ABC model assumes a clear picture of whether a relation exists or not in the literature. For evaluation purposes we opt for a simple criterion to determine the cut-off year: since the discovery pattern tends to show sporadic co-occurences followed by a continuous surge, the cut-off year is defined as the last year with no co-occurrence, i.e. the year just before the relation starts appearing every single year. While not perfect, this criterion appears to match quite well the start of the surge in most cases, as shown by the vertical dashed line in Fig. 1. For the sake of simplicity we arbitrarily pick the year obtained (1) on the “abstracts+articles” view and (2) at the document level. The year is picked independently for each data source (KD and PTC) in order to avoid favouring any of them. In the special case of the FTD-Bipolar relation in PTC no cut-off year can be determined due to the absence of data (see §2.3). Nevertheless this case is not discarded: the relation is counted as “not found” for PTC in order to preserve the comparability of the results.

The methods described below are evaluated on a sample of 8 instances, each made of a target concept and an expected gold-standard related concept. The ideal outcome is that the gold concept is predicted at a position close to the top in the output ranking. However it cannot be assumed that the other concepts related to the target represent a negative relation, therefore performance must be measured in terms of recall only. Additionally the measure must account for the possibility that the gold concept is not retrieved at all by the system, i.e. the rank is undefined. We propose two evaluation measures:

- **R@N**. “Recall at N” is defined as the proportion of instances where the gold concept is found within the top N positions in the ranking (higher is better).
- **MNTR@N**. “Mean Normalized Truncated Rank at N” is defined as the mean across instances of the rank, considering any rank higher or equal to N as N and dividing by N (lower is better because MNTR is essentially an averaged rank).

Example: for three instances a system returns ranks 57, 271 and “not found”. R@100 is 1*/*3 = 0.33. MNTR@100 is calculated as follows: the last two ranks are truncated and replaced with 100; the normalized truncated ranks are 0.57, 1, 1 and the MNTR@100 is the mean of these three values (0.86). R@N is an intuitive but coarse measure, especially with a small sample. MNTR@N provides a more fine-grained evaluation: if two systems return the same number of matches at some threshold N, their R@N is identical but MNTR@N can be used to distinguish them by comparing the average rank. Thus these two measures are complementary: the former provides a “big picture” view of the performance and is easy to interpret, while the latter is slightly less intuitive but provides a more accurate estimation of the performance of a system.

### 3.2. Association measures

Quantifying the strength of association between two concepts in the literature is challenging due to the wide variations in frequency across concepts. Naturally, frequent concepts co-occur with more concepts than rare ones, thus two frequent concepts may co-occur by chance. Rare concepts are usually more specific than frequent ones but they are also harder to distinguish from noise. Similarly to other LBD works (Pyysalo et al., 2019; Crichton et al., 2020), we use the standard Pointwise Mutual Information (PMI) measure as well as some of its variants: Normalized PMI (NPMI) is “slightly less biased than PMI towards low frequency” Bouma (2009); PMI^2^ and PMI^3^ are other alternatives to PMI designed to counter its low frequency bias; [Normalized] Mutual Information ([N]MI) are significantly different in that they take into account all the cases of concepts A and B appearing or not (we follow the definitions from (Bouma, 2009)). We also include Symmetric Conditional Probability (SCP) (Pyysalo et al., 2019), however measures based on raw frequency are excluded because these are strongly biased in favour of concepts which are frequent and generic. Such as bias is inconsistent with our approach meant to extract undiscovered public knowledge (see §2.1), and most of the target relations that we study involve one frequent and one rare concept.

### 3.3. Ranking methods

The contrast method that we propose consists in ranking the concepts in a *reference view* and then removing or downranking the concepts which are highly ranked in a second ranking obtained from a *mask view*. The rationale is that the top concepts in a simple ranking represent the strongest and therefore the most trivial relations, whereas the least associated concepts, which are essentially noise, are found at the bottom of the ranking. Thus the relations of interest which represent relevant undiscovered public knowledge appear at some undetermined rank: after the most trivial relations since their association is not as strong, but before the noise since they are expected to have good association strength. The masking is meant to “hide” the most trivial relations in order to push the relations of interest at the top of the ranking. In this perspective the choice of the reference and mask views is crucial: ideally the reference view is a detailed, broad-coverage and potentially high-noise representation of the literature, while the mask view is a tightly constrained representation supposed to return mostly the trivial relations. To this end we exploit the characteristics of the views presented in §2.2: For example, we expect the full articles to contain all kinds of relations and include noise whereas the abstracts should focus on the essential relations discussed in the paper, making the former a good candidate as reference view and the latter as mask view. Similarly, a view using the level by document can be contrasted with a view by sentence, since the latter is expected to contain only closely related concepts as opposed to the former.

Three contrast methods are tested as well as a baseline method:

- The **baseline** method is a simple ranking of the concepts related to the target. This method can be applied to any data view (see §2.2) and based on any of the association measures. Additionally a minimum and a maximum joint frequency parameters can be applied to filter out some of the concepts, under the assumption that the rarest concepts happen mostly by chance (noise) and the most frequent concepts represent mostly trivial relations.
- In the **basic contrast** method, first the reference view concepts are ranked then the concepts which have a joint frequency in the mask view higher than the maximum parameter are filtered out. This method is similar to using the maximum parameter with the baseline method, except that the filtering is based on the mask view. In other words, the mask view is assumed to represent the strongest (most trivial) relations more reliably than the reference view.
- The **rank difference** contrast method assumes that relations of interest are found across a spectrum where the most relevant concepts are ranked close to the top of the reference view and either absent in the mask view or close the bottom, and conversely. Under this assumption the difference between the rank in the reference view and the one in the mask view is expected to retrieve concepts of interest. Two versions of this method are proposed, one based on the **absolute rank difference** and the other on the **relative rank difference**, where the relative rank is normalized with respect to the size of the view in order to account for the difference in number of concepts between the reference and the mask view.

In the contrast methods the minimum and maximum joint frequency are applied after the masking operation, respectively using the reference and mask view frequency. Additionally an option is added to discard the concepts which appear in the reference view but do not appear at all in the mask view, in the hypothesis that such concepts are mostly noise.

## 4. Results and discussion

### 4.1. Unsupervised experiment

All the experiments presented in this section are run on the set of 8 ND discoveries presented in Table 1, using the cut-off year defined as described in §3.1. In a realistic use case, the concepts would likely be filtered by semantic type in order to target specific categories of relations such as disorders/genes, disorders/drugs or disorders/disorders. Nevertheless concepts are not filtered in these experiments in order to avoid any bias. This implies that the ranks and performance measures shown correspond to a worst-case scenario, since filtering would drastically reduce the search space of the related concepts for every target.

In this first experiment the baseline, basic contrast and absolute difference rank methods are compared across several predefined parameters. The values chosen for the association measure and minimum frequency are arbitrary. The reference and mask view are selected based on the rationale outlined in §3.3. Table 2 shows the MNTR@1000 performance obtained with both the KD and PTC datasets.

**Table 2:**
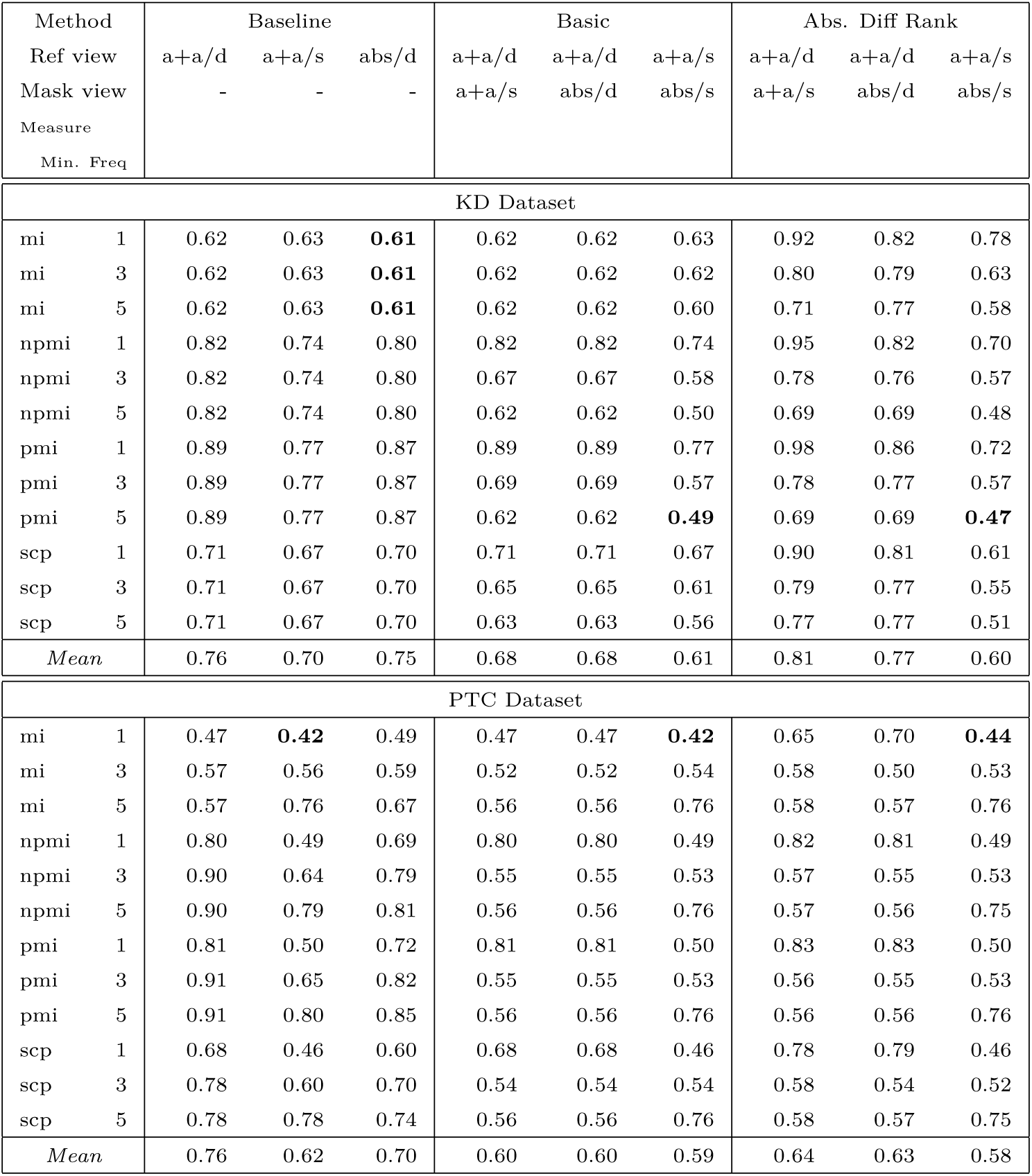
MNTR@1000 performance of three methods depending on measure and minimum frequency for KD and PTC datasets. “a+a” (resp. “abs”) denotes the abstracts+articles (resp. abstracts) view, and “/d” (resp. “/s”) denotes the document (resp. sentence) level. Bold values show the best performance for each method.

| Method |  | Baseline |  |  | Basic |  |  | Abs. Diff Rank |  |  |
| --- | --- | --- | --- | --- | --- | --- | --- | --- | --- | --- |
| Ref view |  | a+a/d | a+a/s | abs/d | a+a/d | a+a/d | a+a/s | a+a/d | a+a/d | a+a/s |
| Mask view |  | - | - | - | a+a/s | abs/d | abs/s | a+a/s | abs/d | abs/s |
| Measure |  |  |  |  |  |  |  |  |  |  |
| Min. Freq |  |  |  |  |  |  |  |  |  |  |
| KD Dataset |  |  |  |  |  |  |  |  |  |  |
| mi | 1 | 0.62 | 0.63 | **0.61** | 0.62 | 0.62 | 0.63 | 0.92 | 0.82 | 0.78 |
| mi | 3 | 0.62 | 0.63 | **0.61** | 0.62 | 0.62 | 0.62 | 0.80 | 0.79 | 0.63 |
| mi | 5 | 0.62 | 0.63 | **0.61** | 0.62 | 0.62 | 0.60 | 0.71 | 0.77 | 0.58 |
| npmi | 1 | 0.82 | 0.74 | 0.80 | 0.82 | 0.82 | 0.74 | 0.95 | 0.82 | 0.70 |
| npmi | 3 | 0.82 | 0.74 | 0.80 | 0.67 | 0.67 | 0.58 | 0.78 | 0.76 | 0.57 |
| npmi | 5 | 0.82 | 0.74 | 0.80 | 0.62 | 0.62 | 0.50 | 0.69 | 0.69 | 0.48 |
| pmi | 1 | 0.89 | 0.77 | 0.87 | 0.89 | 0.89 | 0.77 | 0.98 | 0.86 | 0.72 |
| pmi | 3 | 0.89 | 0.77 | 0.87 | 0.69 | 0.69 | 0.57 | 0.78 | 0.77 | 0.57 |
| pmi | 5 | 0.89 | 0.77 | 0.87 | 0.62 | 0.62 | **0.49** | 0.69 | 0.69 | **0.47** |
| scp | 1 | 0.71 | 0.67 | 0.70 | 0.71 | 0.71 | 0.67 | 0.90 | 0.81 | 0.61 |
| scp | 3 | 0.71 | 0.67 | 0.70 | 0.65 | 0.65 | 0.61 | 0.79 | 0.77 | 0.55 |
| scp | 5 | 0.71 | 0.67 | 0.70 | 0.63 | 0.63 | 0.56 | 0.77 | 0.77 | 0.51 |
| Mean |  | 0.76 | 0.70 | 0.75 | 0.68 | 0.68 | 0.61 | 0.81 | 0.77 | 0.60 |
| PTC Dataset |  |  |  |  |  |  |  |  |  |  |
| mi | 1 | 0.47 | **0.42** | 0.49 | 0.47 | 0.47 | **0.42** | 0.65 | 0.70 | **0.44** |
| mi | 3 | 0.57 | 0.56 | 0.59 | 0.52 | 0.52 | 0.54 | 0.58 | 0.50 | 0.53 |
| mi | 5 | 0.57 | 0.76 | 0.67 | 0.56 | 0.56 | 0.76 | 0.58 | 0.57 | 0.76 |
| npmi | 1 | 0.80 | 0.49 | 0.69 | 0.80 | 0.80 | 0.49 | 0.82 | 0.81 | 0.49 |
| npmi | 3 | 0.90 | 0.64 | 0.79 | 0.55 | 0.55 | 0.53 | 0.57 | 0.55 | 0.53 |
| npmi | 5 | 0.90 | 0.79 | 0.81 | 0.56 | 0.56 | 0.76 | 0.57 | 0.56 | 0.75 |
| pmi | 1 | 0.81 | 0.50 | 0.72 | 0.81 | 0.81 | 0.50 | 0.83 | 0.83 | 0.50 |
| pmi | 3 | 0.91 | 0.65 | 0.82 | 0.55 | 0.55 | 0.53 | 0.56 | 0.55 | 0.53 |
| pmi | 5 | 0.91 | 0.80 | 0.85 | 0.56 | 0.56 | 0.76 | 0.56 | 0.56 | 0.76 |
| scp | 1 | 0.68 | 0.46 | 0.60 | 0.68 | 0.68 | 0.46 | 0.78 | 0.79 | 0.46 |
| scp | 3 | 0.78 | 0.60 | 0.70 | 0.54 | 0.54 | 0.54 | 0.58 | 0.54 | 0.52 |
| scp | 5 | 0.78 | 0.78 | 0.74 | 0.56 | 0.56 | 0.76 | 0.58 | 0.57 | 0.75 |
| Mean |  | 0.76 | 0.62 | 0.70 | 0.60 | 0.60 | 0.59 | 0.64 | 0.63 | 0.58 |

First, important variations in the performance between the two data sources can be observed. In particular the frequency threshold appears to have a different effect with the contrast methods: while increasing the threshold almost always increases performance with KD, the effect is not constant in PTC; a minimum frequency of 5 gives the best average performance in KD but the worst in PTC. This is likely due to the fact that the KD and PTC processes differ and do not always extract the same concepts (see also §2.3). Similarly the baseline method is quite sensitive to the frequency threshold in PTC but not in KD. However the comparison of the mean performance between the different methods shows a roughly similar pattern: the basic contrast method outperforms the baseline by a significant margin, with the rank difference method performance between the two other methods.

Detailed observation of the results obtained by the different association measures shows a pattern found in both KD and PTC: in average MI obtains the best results followed by SCP, whereas PMI and NPMI perfom significantly worse. The specific parameters of the contrast methods also have an important impact on performance, with the contrast between the *abstracts and articles* view masked by the *abstracts* view at sentence level performing the best in average for both data sources, albeit with a smaller difference in PTC.

### 4.2. Supervised Experiment

The contrast method and even the baseline method are *parametric heuristics*: the method itself is unsupervised and does not require any fitting to the data, however it accepts several parameters which impact the output (see §3). The unsupervised experiment above provides an approximate evaluation of the performance of the method in its main use case, i.e. when used by a human who would select what they think are reasonable values for the parameters. However the selection of the parameters is arbitrary, therefore the performance estimation cannot be considered statistically reliable. In the next experiment we propose a different use case where the parameters are tuned during a training stage. This use case corresponds to a data-driven approach where the system would be trained based on a dataset consisting of a decent number of known discoveries. The discoveries in this training set should preferably be recent and chosen to represent the type of relation that one wants to retrieve. To some extent this setting is similar to the quantitative approach used for example by Lever et al. (2017) (see §2.1), and allows a more reliable estimation of the performance of the methods.

In this experiment we use leave-one-out cross-validation, which is roughly equivalent to 10-fold cross-validation given the small side of our dataset (8 relations). At every iteration a large set of parameters settings are applied to the target concepts in the training set, then the performance is evaluated across the training set relations. The parameter setting which reaches the best performance on the training set is then selected and applied to the test instance. Finally the performance is calculated across the test instances. This parameter tuning process corresponds to an exhaustive search, with the parameters ranging all the possible values.^11^ A total of 52,080 combinations of parameters are tested. Given the small size of the training set (only 7 cases) there is a risk of overfitting. While the variance is very high, there was no indication of overfitting when comparing the performance between the training and test set. We hypothesize that the variance is caused not only by the small size of the training set but also by the important differences between the different cases (different cut-off years, frequency patterns, etc.).

Table 3 shows the performance obtained by the different methods. These results confirm that the basic contrast method performs best with both data sources, but not by a large margin. It also shows that the baseline performs better than any of the two rank difference methods. The MNTR@1000 performance is not very different from the means obtained in Table 2. In particular the performance obtained with supervised tuning does not reach the best cases observed with arbitrary parameters, wich indicates that these cases certainly happened by chance. In general better performance is obtained using the KD source, although the R@500 values are higher with PTC. This would suggest that target relations tend to be ranked either close to the top or far down the ranking with PTC, but the sample is too small to draw this conclusion.

**Table 3:** Performance of the methods in the supervised experiment.

| Data source | Method | R@N |  |  | MNTR@N |  |  |
| --- | --- | --- | --- | --- | --- | --- | --- |
|  |  | 100 | 500 | 1000 | 100 | 500 | 1000 |
| KD | Baseline | 0.12 | 0.38 | 0.62 | 0.93 | 0.72 | 0.63 |
| KD | Basic Contrast | 0.25 | 0.50 | 0.62 | 0.83 | 0.64 | 0.54 |
| KD | Abs. Diff Rank | 0.25 | 0.38 | 0.62 | 0.90 | 0.71 | 0.60 |
| KD | Rel. Diff Rank | 0.25 | 0.38 | 0.62 | 0.85 | 0.71 | 0.56 |
| PTC | Baseline | 0.00 | 0.50 | 0.50 | 1.00 | 0.76 | 0.63 |
| PTC | Basic Contrast | 0.12 | 0.50 | 0.50 | 0.99 | 0.72 | 0.61 |
| PTC | Abs. Diff Rank | 0.00 | 0.38 | 0.50 | 1.00 | 0.83 | 0.67 |
| PTC | Rel. Diff Rank | 0.12 | 0.50 | 0.50 | 0.99 | 0.79 | 0.65 |

By observing how often a particular value is selected as the best option, this experiment also offers some additional insight about the parameters. Surprisingly in several cases the optimal value of the parameter appears to primarily depend on the data source, as shown in Table 4. This is expected in the case of the minimum frequency, which is optimal at a very low level (1 or 2) in PTC but at a high level (7 to 9) in KD. But the optimal association measure appears to strongly depend on the data source as well: MI is selected in 64% of the cases, whereas PMI is the most selected in KD (NMI and PMI^3^ are also frequently selected). The selection of the reference/mask views shows the sharpest difference between data sources:

- In the optimal combination for PTC, the reference view is almost always *abstracts+articles* at document level and the mask is the *abstracts* view at either document or sentence level. This is consistent with our rationale (see §3.3): the reference is made of a large volume of possibly noisy relations, while the mask view contains a stricter subset of more reliable relations.
- However for KD the optimal combinations appear to counter our intuition: the most frequent selected combination is made of abstracts as reference view and abstracts+articles as mask, both at document level.

**Table 4:**
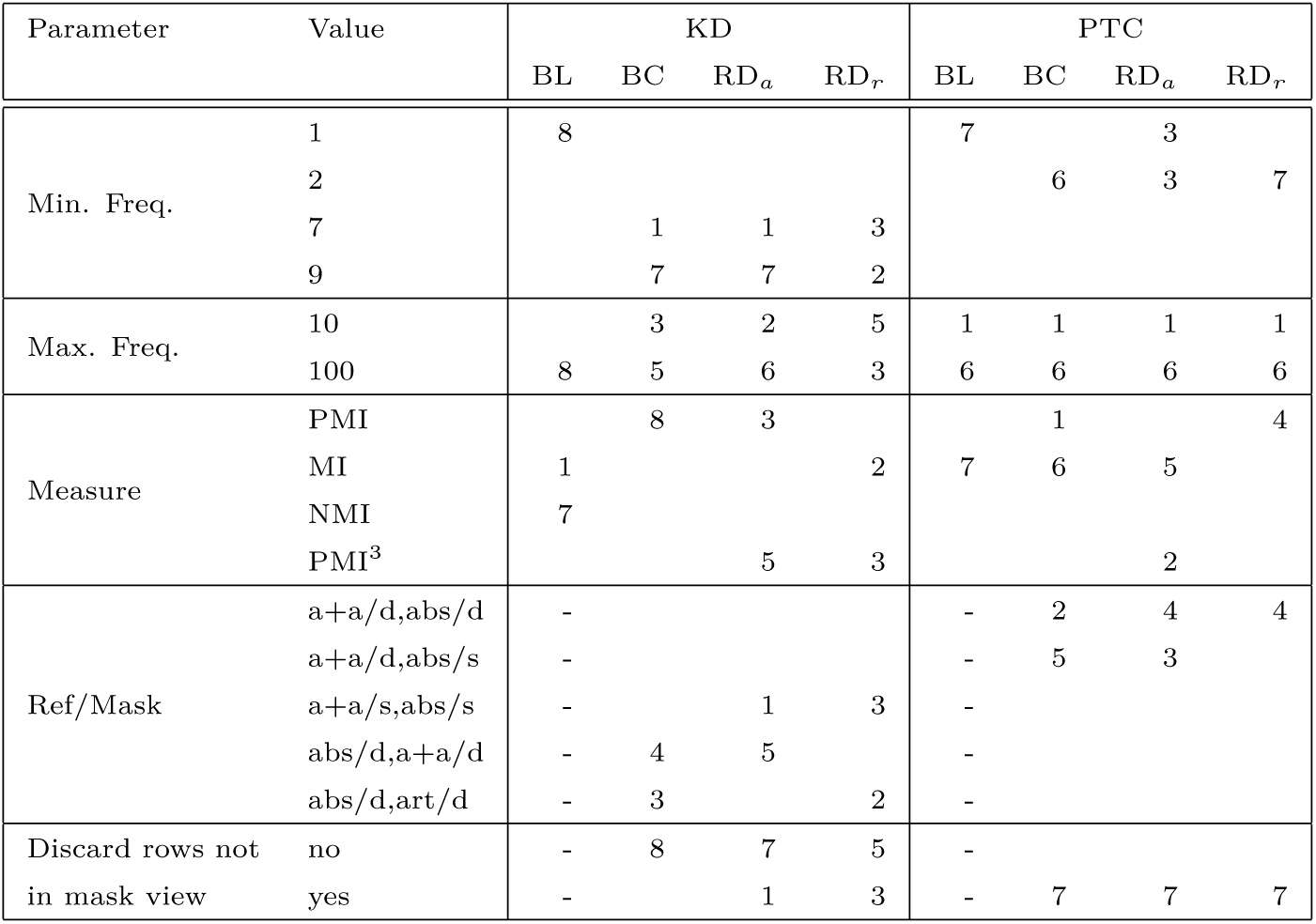
Frequency of the parameters values selected as optimal in the supervised experiment. BL, BC, RD_*a*_ and RD_*r*_ stand respectively for baseline, basic contrast, absolute/relative rank difference. In the pairs of reference and mask view, “a+a” (resp. “abs”) denotes the abstracts+articles (resp. abstracts) view, and “/d” (resp. “/s”) denotes the document (resp. sentence) level. Rare values (selected less than 3 times) are not shown.

Additionally the option to remove the concepts absent in the mask view is consistently selected with PTC but rarely with KD. It is difficult to interpret these differences with precision, but it is clear that differences in the concepts extraction and disambiguation process cause serious discrepancies in the knowledge base.

Overall this last experiment shows a modest performance improvement of the basic contrast method over the baseline method, but it should be noted that the baseline itself is fairly competitive since its parameters are tuned similarly to the contrast methods.

### 4.3. Online exploration tool

The contrast method is intended to be used by medical experts through a simple, user-friendly exploration tool. A simple prototype has been developed and is available publicly at brainmend.adaptcentre.ie. The use case is the retrieval of potentially relevant relations from the full existing literature. The method itself is meant to be transparent and intuitive to the user, and the exploration tool is designed accordingly. It is made of two parts:

- The first part called “Top associated concepts” lets the user select a target concept among a predefined list and shows the list of related concepts ordered by association strength. The user can select the various parameters and filter the output concepts by semantic type. In particular they can experience the difference in the results displayed depending on the selected view (see §2.2).
- The second part called “Contrast two datasets” applies the contrast method to two data views selected by the user. The user can interact with the parameters until they obtain the desired type of result as output. In particular the user can use the minimum and maximum frequency thresholds in order to tune the level of specificity/genericity in the output concepts: it is fairly intuitive that the higher the frequency the more generic the concepts, and the user can observe the effect of their actions immediately.

The contrast method relies on the availability of different “views” of the literature. For the sake of usability, in the prototype the rankings for every target and every view must have been preprocessed offline. This is a disadvantage since in the current implementation the user must choose among a small set of predefined targets.

## 5. Conclusion

Through this work we hope to encourage the development of new LBD methods beyond the traditional ABC paradigm. Despite its intuitive appeal, there is no evidence that the ABC model can adequately cover the diverse types of discoveries found in the literature, as demonstrated by our research. We proposed the contrast approach for LBD which achieves a good performance on our data, but further experimental validation is needed given the high variance across cases. In this work we also uncovered serious discrepancies between data sources, an indication that LBD methods would likely benefit from diversifying their sources instead of relying on a single one.

## Acknowledgements

This work was conducted with the financial support of Irish Health Research Board (HRB) BRAINMend project (grant number HRB-JPI-JPND-2017-1) and with the support of the Science Foundation Ireland Research Centre ADAPT (grant number 13/RC/2106_P2).

## Footnotes

1 It is worth noting that the ABC model was introduced in 1986: at the time ML-based approaches were in their infancy and rule-based systems were still a significant part of the state of the art in information retrieval.

2 For both sources the data used in the experiments was downloaded in January 2021.

3 https://www.nlm.nih.gov/databases/download/pubmed_medline.html

4 https://www.ncbi.nlm.nih.gov/pmc/tools/openftlist/

5 https://github.com/erwanm/knowledgediscovery.

6 https://www.nlm.nih.gov/research/umls/index.html

7 https://github.com/erwanm/kd-data-tools.

8 https://www.ncbi.nlm.nih.gov/research/pubtator/

9 https://www.ncbi.nlm.nih.gov/mesh. MeSH stands for Medical Subjects Headings.

10 The case Parkinson-ALS is omitted in order to preserve readability because the scale of the frequencies is much higher than in the other cases. The years 2019 and 2020 are omitted because they are not complete yet in the data source.

11 The minimum frequency is restricted to the range [1,10] and the maximum to only 4 values: 10,100, 1000, NA. NA means that no maximum threshold is applied.

